# Enhancement of Arabidopsis growth by *Enterobacter* sp. SA187 under elevated CO_2_ is dependent on ethylene signalling activation and primary metabolism reprogramming

**DOI:** 10.1101/2025.07.08.663752

**Authors:** Amina Ilyas, Caroline Mauve, Stéphanie Pateyron, Christine Paysant-Le Roux, Jean Bigeard, Michael Hodges, Axel de Zélicourt

## Abstract

As atmospheric CO_2_ levels continue to increase, optimizing the CO_2_ fertilization effect which often falls short of its potential due to the physiological and metabolic limitations of plants becomes crucial. This study investigates the role of *Enterobacter* sp. SA187 (SA187), a plant growth-promoting bacterium, in enhancing growth and development of *Arabidopsis thaliana* under elevated atmospheric CO_2_ (eCO_2_) conditions. While SA187 inoculation did not have major effects under ambient CO_2_, it was found to significantly enhance root and shoot biomass, and to increase N- and reduce C-contents under eCO_2_. Moreover, transcriptomics and metabolomics suggested that SA187 modulated phytohormonal homeostasis, with activation of the salicylic acid, jasmonic acid and ethylene signalling pathways, and increased primary metabolism including the TCA cycle, N and carbohydrate metabolisms. Finally, the growth-promoting effects of SA187 were shown to be mediated through ethylene-dependent pathways, as evidenced with the ethylene-insensitive mutant *ein2-1* which did not show similar benefits in plant fresh weight and altered gene expression. This beneficial plant-microbe interaction under eCO_2_ in a non-leguminous plant highlights a novel aspect of microbial influence on plant physiology in the context of climate change. These insights underscore the potential of utilizing SA187 to enhance plant performance and adaptability in future high CO_2_ environments, providing a sustainable approach to agricultural productivity as global CO_2_ levels increase.

## I. Introduction

The hallmark of the Anthropocene is the rise in atmospheric CO_2_ levels, which have increased from approximately 280 ppm in the 17th century to over 430 ppm today (NOAA, 2025). Projections suggest that without significant mitigation efforts, these levels could reach between 800 to 1000 ppm by the end of the century (Breecker *et al*., 2010; Meinshausen *et al*., 2011). Moreover, the accumulation of greenhouse gases is driving climate change, altering global weather patterns, leading to unpredictable seasons and shifts in temperature and precipitation (Abbass *et al*., 2022; Mikhaylov *et al*., 2020).

Elevated atmospheric CO_2_ concentration (eCO_2_) has a direct positive effect on plant growth, development and productivity, by enhancing photosynthetic CO_2_ assimilation and lowering wasteful photorespiration. However, this eCO_2_ fertilization effect is often lower than predicted and it can depend on plant species and environmental factors (Rho *et al*., 2020; Ainsworth and Long, 2021). The lower-than-expected increase is the consequence of plant photosynthetic acclimation during long-term eCO_2_ exposures, which is attributed to various physiological and metabolic factors (Long *et al*., 2004; Ainsworth and Long, 2005; Leakey et al, 2009). Plants growing in an eCO_2_ atmosphere also exhibit a reduction in plant tissue mineral nutrient concentrations that adversely affect crop quality (Gojon *et al*., 2022). An average decrease in N-content of 10-15 % is often observed in eCO_2_ grown plants although this is highly variable (Taub and Wang, 2008). These eCO_2_ acclimations are associated with a complex reprogramming of gene expression (Moore *et al*., 1999; Cheng *et al*., 1998), leading to lower amounts of ribulose-1,5-bisphosphate carboxylase/oxygenase (Rubisco) protein, and a reduction in nitrate uptake and assimilation (Rubio-Asensio *et al*., 2017). Photosynthetic acclimation to eCO_2_ is associated with sink limitations, leading to an accumulation of non-structural carbohydrates in photosynthetic tissues (Taub and Wang, 2008; Aranjuelo *et al*., 2011; Rogers and Ainsworth, 2006) and hexokinase-dependent sugar signalling (Cheng *et al* 1998; Moore *et al* 1999; Peyo *et al* 2000; Long *et al* 2004) that negatively impact, among others, photosynthetic gene expression ( Krapp *et al*., 1993; Moore *et al*., 1999) and thus photosynthetic activity (Pérez *et al*., 2011; Aranjuelo *et al*., 2013). Improved sink strength has been associated to a reduction in eCO_2_ acclimation as reported in tobacco (Ruiz-Vera *et al* 2017), rice (Zhu *et al* 2014) and barley (Torralbo *et al* 2019).

Several factors have been associated with the reduction of N content under eCO_2_. Much of the decrease can be accounted for by the reduction in Rubisco protein content, a finding supported by Free Air Concentration Enrichment (FACE) studies (Ainsworth & Long, 2005). It is also believed that eCO_2_ leads to the dilution of N in plant tissues due to an increased and faster growth (Taub *et al*., 2008; Taub & Wang, 2008; Gifford *et al*. 2000, Gojon *et al*., 2023). A lowering of leaf transpiration from eCO_2_-related changes in both stomatal density and aperture size can reduce N-uptake and alter N-partitioning (McDonald *et al*. 2002; Long *et al*., 2004; Ainsworth and Rogers, 2007). It has also been proposed that nitrate uptake and subsequent shoot nitrate assimilation are both negatively impacted by eCO_2_-associated inhibition of photorespiration (Rachmilevitch *et al*., 2004). Therefore, it appears that maintaining a balance between photosynthetic CO_2_ assimilation and N uptake and assimilation is crucial to sustain the eCO_2_ fertilization effect and this is reflected in the plant C/N ratio (Ruiz-Vera *et al*., 2017, Kant *et al*., 2012).

A strategy to enhance plant resilience to reduce the negative effects of eCO_2_ acclimations could be the exploitation of plant beneficial microbes. Plant growth-promoting bacteria (PGPB) have received particular attention due to their diversity, abundance, and ease of manipulation (Glick, 2012). PGPB have been shown to improve plant growth and development under normal and various abiotic stress conditions, such as drought, salinity, and heat, by modulating plant hormonal balance including auxins, ethylene, cytokinins and gibberellins (Persello-Cartieaux *et al*., 2003; Vessey, 2003; Hardoin *et al*., 2008), improving root architecture, and enhancing mineral acquisition by N-fixation, phosphate and zinc solubilisation, and iron sequestration (Compant *et al*., 2010). PGPB can also produce antimicrobial agents and induce systemic resistance against plant pathogens (Glick, 2012; Pieterse *et al*., 2008).

In the context of eCO_2_, it has been shown that the symbiotic relationship between leguminous plant species and N_2_-fixing rhizobia located in root nodules appear to be less affected by eCO_2_ acclimations when compared to many non-leguminous plants (Cotrufo *et al*. 1998; Jablonski *et al*. 2002; Taub *et al*. 2008). This is believed to be due to rhizobia consuming large quantities of plant photosynthates in exchange for fixed N, thus providing a strong carbon-sink and reducing N-limitations (Irigoyen *et al*., 2014; Singer *et al*., 2020). However, there is no report of PGPB maintaining an eCO_2._fertilization effect in non-leguminous C3 plants.

A well-studied PGPB shown to improve plant resilience to climate change-associated abiotic stresses such as salt or heat is *Enterobacter* sp. SA187 (hereafter named SA187) (de Zélicourt *et al*., 2018; Shekhawat *et al*., 2021). This endophytic bacterium was isolated from root nodules of *Indigofera argentea*, a leguminous plant found in the Saudi Arabian desert (Andres Barrao, et al 2017). In *Arabidopsis thaliana* it was shown to colonize the surface and inner tissues of both roots and shoots (de Zélicourt *et al*., 2018). The functional analysis of the SA187 genome indicated genes involved in nutrient uptake/exchange, chemotaxis, plant colonization, and oxidative stress. Moreover, bacterial metabolic pathways were identified that could potentially contribute to plant growth promotion (Andres Barrao, et al 2017).

SA187 improved the abiotic stress tolerance of *in vitro* grown *Arabidopsis thaliana* and field-grown *Medicago sativa* (alfalfa) and *Triticum durum* (wheat) (de Zélicourt *et al*., 2018; Shekhawat *et al*., 2021). This was in part due to the maintenance of primary metabolism including photosynthesis (de Zélicourt *et al*., 2018). Ethylene sensing was found also to play an important role since Arabidopsis mutants impaired in ethylene perception no longer exhibited a beneficial response to SA187 while this was maintained in ethylene synthesis mutants (de Zélicourt 2018, Shekhawat *et al*., 2021).

This study aims to explore whether SA187 can enhance plant growth under eCO_2_ conditions. Based on transcriptomic and metabolomic analyses, initial findings indicate that SA187 stimulates growth under eCO_2_ through the activation of several phytohormonal signalling pathways while modulating plant N and carbohydrate metabolisms including the TCA cycle. This research highlights how plant-microbial interactions can be harnessed to support C3 plant acclimations to rapidly changing global climates, offering new perspectives to help sustainable agricultural practices.

## II. Materials and methods

### A. Bacterial and plant materials

*Enterobacter sp.* SA187 was originally isolated from root nodules of *Indigofera argentea* in Jizan, Saudi Arabia (Andrés-Barrao *et al*., 2017). *Arabidopsis thaliana* ecotype Columbia-0 (Col-0, wild-type) and the ethylene-insensitive mutant *ein2-1* (Guzman *et al*., 1990) were cultivated under controlled greenhouse conditions (16 h photoperiod, 18/20°C night/day, 50 ± 10% relative humidity) for seed propagation.

### B. Plant growth and bacterial colonization

Seed sterilization and bacterial inoculation were performed as described previously (Saad *et al*., 2018). Briefly, sterilized Arabidopsis seeds were sown on half-strength Murashige and Skoog (½ MS) medium (Duchefa basal salts only, 0.5 g/L MES, pH 5.8, 9 g/L agar) containing SA187 (+SA187, concentration = 2.10^5^ cells/mL) or not (mock) and stratified for 24 h in the dark at 4°C. These plates were then placed vertically in growth chambers for seed germination (16 h photoperiod, 18/20°C night/day, 50±10% relative humidity), either at 450 ppm CO_2_ for aCO_2_ conditions or 1000 ppm CO_2_ for eCO_2_ conditions, resulting in 4 experimental groups: (i) Mock_aCO_2_ (ii) SA187_ aCO_2_, (iii) Mock_eCO_2_, (iv) SA187_eCO_2_.

At 5 days post-germination (dpg), seedlings were transferred to fresh ½ MS agar plates and grown vertically. Primary root length (PRL) was measured using ImageJ software after scanning of the plates 12 dpg. Lateral root density (LRD) was evaluated as detectable number of lateral roots under a stereo microscope divided by the PRL at 12 dpg. Fresh weight (FW) of shoots and roots was measured 15 dpg of seedlings. Dry weight (DW) was measured after drying shoot and roots for 2 days at 70°C. Bacterial colonization was quantified as described previously (Saad *et al*., 2018). Samples were immediately flash-frozen in liquid-N_2_ and stored at –80°C for subsequent analyses.

### C. Transcriptomics

#### 1. RNA-sequencing and data analyses

Shoot and root samples from 15 dpg plants from all experimental groups at a 1.03 developmental growth stage were collected for RNAseq experiments to obtain 3 biological replicates (Boyes, 2001). Each replicate was composed of either rosettes or roots from 4 to 6 plants per condition. Total RNA was extracted using Nucleospin RNAplus kit (Macherey Nagel), according to the supplier’s instructions and further purified using the RNA Clean & Concentrator Kits (Zymo Research®, California, USA). RNA-seq libraries were constructed using the TruSeq Stranded mRNA library prep kit (Illumina®, California, USA) according to the supplier’s instructions. Libraries were sequenced in single-end (SE) mode with 75 bases for each read on a NextSeq500 to generate between 16 and 45 millions of reads per sample.

Adapter sequences and bases with a Q-Score below 20 were trimmed out from reads using Trimmomatic (v0.36, Bolger *et al*. 2014) and reads shorter than 30 bases after trimming were discarded. Reads corresponding to rRNA sequences were removed using sortMeRNA (v2.1, Kopylova E. *et al*. 2012) against the silva-bac-16s-id90, silva-bac-23s-id98, silva-euk-18s-id95 and silva-euk-28s-id98 databases.

Filtered reads were then mapped and counted using STAR (v2.7.3a, Dobin *et al*. 2013) with the following parameters --alignIntronMin 5 --alignIntronMax 60000 -- outSAMprimaryFlag AllBestScore --outFilterMultimapScoreRange 0 -- outFilterMultimapNmax 20 on the *Arabidopsis thaliana* genome (ARAPORT version 11) and its associated GTF annotation file. At least 97% of the reads were associated to annotated genes.

Statistical analyses were carried out separately on shoots and roots with R v3.6.2 (R Core Team, 2020) using the Bioconductor package edgeR (v 3.28.0, Robinson *et al*., 2010; McCarthy *et al*., 2012). For both analyses, low counts genes were filtered using the “filterByExpr” function of the R package edgeR with a minimum count threshold equal to 15. Raw counts were normalized using the trimmed mean of M values (TMM) method. Differential analyses were based on a negative binomial generalized linear model. For each analysis, the log2 of the average normalized gene expression is an additive function of a CO_2_ condition factor (2 modalities), a treatment factor (2 modalities), a replicate factor and an interaction between the CO_2_ condition factor and the treatment factor. In each analysis, by using a likelihood ratio test, the difference between the two treatments at each CO_2_ condition, the difference between the two CO_2_ conditions given a treatment and the interaction effect defined as the difference between the two treatments at eCO_2_ minus the difference between the same two treatments at ambient CO_2_ were evaluated. The distribution of the raw p-values were checked following the quality criterion described by Rigaill et al., 2018 and a gene was declared differentially expressed (DEG) if its adjusted p-value was lower than 0.05. Hierarchical clustering was done using MeV software and GO-term enrichment analysis was performed with g:Profiler (Raudvere *et al*., 2019).

#### 2. qRT-PCR analyses

Extracted mRNA (see above) was reverse-transcribed using the Promega ImProm-II™ Reverse Transcription System with oligo-dT primers, following the manufacturer protocol. The resulting cDNA was used for qRT-PCR. Reactions were performed and carried out with a LightCycler® 480 SYBR Green I Master mix (Roche) and the following thermal cycling conditions: A pre-incubation at 95°C for 10 min; 40 cycles of amplification (95°C for 10 s, 60 °C for 10 s, and 72 °C for 10 s); and a dissociation step (melting curve) to validate the PCR products. Gene expression was analyzed using the ΔΔCt method (Rao *et al*., 2013), normalized against two constitutive reference genes (*Actin* and *YSL8*). Primer sequences are provided in Supplementary Material (Sup Table 1).

### D. Metabolomics

#### 1. Extraction

Freeze-ground samples (50 mg FW) were resuspended in water/acetonitrile/isopropanol (2:3:3) with 4 µg/mL ribitol (internal standard), shaken (4°C, 10 min), centrifuged (16000 g, 15 min), and dried (30°C, 4 h) in a Speed-Vac and stored at −80°C.

#### 2. GC-MS analyses

All steps were carried out as described previously (Fiehn, 2006; Fiehn *et al*., 2008). Dry aliquot of 150 µl of the Extraction solution were taken and dried for a second time in a Speed-Vac evaporator for 2 h at 30°C at 14000 rpm before adding 10 µL of 20 mg.mL^-1^ methoxyamine in pyridine to the samples. The first step of derivatization was performed for 90 min at 30°C under continuous shaking in an Eppendorf thermomixer. Then 90 µL N-methyl-N-trimethylsilyl-trifluoroacetamide (MSTFA) (Regis Technologies, Morton Grove, IL, USA) were added and the reaction was continued for 30 min at 37°C. After cooling, all samples were transferred to an Agilent vial for injection.

One microliter of each sample was injected into an Agilent 7890B gas chromatograph coupled to an Agilent 5977A mass spectrometer. The column was an Rxi-5SilMS from Restek (30 m with 10 m Integra-Guard column). An injection in split mode with a ratio of 1:30 was systematically performed for saturated compound quantification. Oven temperature ramp was 60°C for 1 min then 10°C min^-1^ to 325°C for 10 min. Helium constant flow was 1.1 mL.min^-1^. Temperatures were the following: injector: 250°C, transfer line: 290°C, source: 230°C and quadrupole: 150°C. The quadrupole mass spectrometer was switched on after a 5.90 min solvent delay time, scanning from 50 to 600 m/z. Absolute retention times were locked to the internal standard Ribitol using the RTL system provided by Agilent’s Masshunter software. The Agilent Fiehn GC/MS Metabolomics RTL Library (version June 2008) was employed for metabolite identifications. Peak areas were determined using the Masshunter Quantitative Analysis (Agilent Technologies, Santa Clara, CA, USA) in splitless and split 30 modes. Resulting areas were compiled into one single MS Excel file for comparisons. Peak areas were normalized to Ribitol and DW. A total of 114 metabolites were identified and expressed in arbitrary units (semi-quantitative determination). Pathway enrichment analysis was done using MetaboAnalyst 6.0 (Pang *et al*., 2024).

#### 3. Total C and N contents

Lyophilized shoot and root samples (1 mg) were combusted in a Pyrocube Elemental Analyzer (Elementar, France), and N_2_/CO_2_ levels were quantified using a thermal conductivity detector (TCD) against standards (ammonium sulfate 21.2% N, benzoic acid 68.85% C, glutamic acid 9.51% N / 40.82% C and glutamine 19.17% N / 41.09% C). Elemental C and N contents are given in % (mass fraction).

## III. Results

### A. SA187 increases plant growth and development under eCO_2_

To evaluate the impact of SA187 on the growth and development of *Arabidopsis thaliana*, plants were cultivated under aCO_2_ (450 ppm) or eCO_2_ (1000 ppm) conditions, with or without SA187 inoculation (+SA187 or mock), following previously established protocols (de Zélicourt *et al*., 2018; Saad *et al*., 2018). Growth parameters, including FW, DW, PRL, and LRD were assessed at 10 dpg (Figure 1).

**Figure 1:**
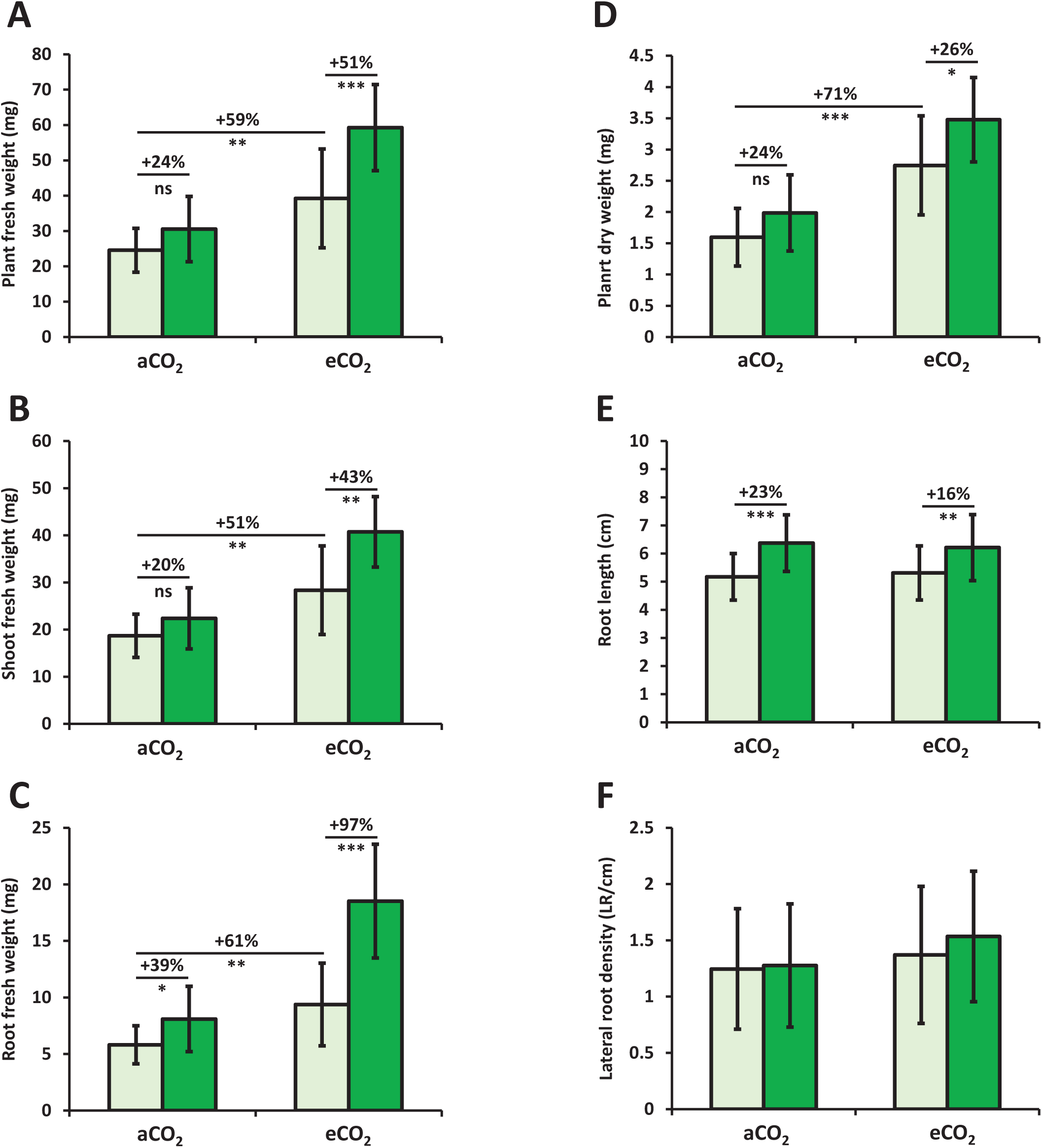
Growth phenotyping of Arabidopsis *Col-0* grown under aCO_2_ and elevated eCO_2_ conditions. **(A)** Plant fresh weight, **(B)** Shoot fresh weight, **(C)** Root fresh weight and **(D)** Total dry weight of plant 15 dpg on ½ MS medium under ambient (aCO_2_) and elevated CO_2_ (eCO_2_) without (Mock) and with (SA187) SA187. **(E)** Primary root length and **(F)** Lateral root density of plants 12 dpg under ambient (aCO_2_) and elevated CO_2_ (eCO_2_) without (Mock) and with (SA187) SA187. Light green indicate mock plants and dark green SA187 inoculated plants. For each measured parameter and every condition, data represent means (n > 30, 3 independent experiments) with standard deviations. Asterisks indicate a statistical difference based on two-sided, unpaired Student’s t-test: \**p* < 0.05; \*\**p* < 0.01; \*\*\**p* < 0.001; ns, not significant.

In uninoculated plants, as expected, eCO_2_ significantly increased plant biomass, with total FW and DW increasing by 59% and 71%, respectively, compared to plants grown under aCO_2_ (Figure 1A). This growth enhancement affected equally roots and shoots with FW increases of 61% and 51% respectively (Figure 1A). These results align with past studies on CO_2_ fertilization in C3 plants, which report enhanced biomass accumulation under eCO_2_ (Ainsworth & Long, 2005, Ainsworth and Long, 2020).

Under aCO_2_ conditions, SA187 did not significantly impact overall plant biomass (Figure 1 A and D), except for a 39% increase in root FW and a 23% increase in PRL (Figure 1 C and E), indicating an effect on root development (Figure 1C). However, under eCO_2_, SA187 markedly improved the total FW (+51%), shoot FW (+43%), root FW (+97%), total DW (+26%) compared to mock plants grown under eCO_2_ conditions (Figure 1, A-D). These results demonstrated that SA187 could significantly enhance the growth and development of *Arabidopsis thaliana* under eCO_2_ conditions. In addition, SA187 increased PRL by a similar magnitude under both aCO_2_ (+23.2%) and eCO_2_ (+16.9%), suggesting that SA187 promotes primary root elongation independently of atmospheric CO_2_ levels. In contrast, SA187 did not influence LRD under either CO_2_ condition, implying that its beneficial effect is specific to primary root growth rather than lateral root formation (Figure 1E-F).

As expected, the well-documented fertilization effect of eCO_2_ was observed. Interestingly, while SA187 did not exhibit any significant beneficial effect on plant biomass under aCO_2_ conditions, with the exception of root FW, it noticeably increased various growth parameters under eCO_2_. The significant increase in plant FW under eCO_2_ conditions, which was augmented further in the presence of SA187, suggests a synergistic interaction between SA187 and eCO_2_ in promoting biomass accumulation. Such a synergy aligns with previous reports demonstrating that SA187 enhances plant biomass and stress tolerance under various abiotic conditions, including salinity and heat stress (de Zélicourt *et al*., 2018). Moreover, SA187 increased PRL under both aCO_2_ and eCO_2_ conditions, but did not influence LRD. In the literature, SA187 was found to increase LRD under salt stress conditions, indicating that the effect of SA187 on root architecture may be context-dependent (de Zélicourt *et al*., 2018). This suggests that SA187 can modulate different regulatory pathways according to environmental context.

Overall, these results highlight SA187’s ability to enhance further plant growth under eCO_2_ when compared to the non-inoculated plants perhaps by reducing acclimation processes that limit the eCO_2_ fertilization effect. The observed synergy between SA187 and eCO_2_ revealed the bacterium’s capability to modulate plant physiological processes to optimize root and shoot growth, reinforcing its value as a biostimulant for enhancing plant development in the face of escalating atmospheric CO_2_ levels.

### B. eCO_2_ does not benefit SA187 proliferation and plant colonization

To investigate whether the enhanced plant growth effects in the presence of SA187 under eCO_2_ were associated with changes in bacterial proliferation, SA187 was cultured in liquid medium under both aCO_2_ and eCO_2_ conditions. Optical density measurements taken over 10 h showed no significant differences in bacterial proliferation between the two conditions (Figure 2A), indicating that eCO_2_ does not directly influence SA187 proliferation *in vitro*.

**Figure 2:**
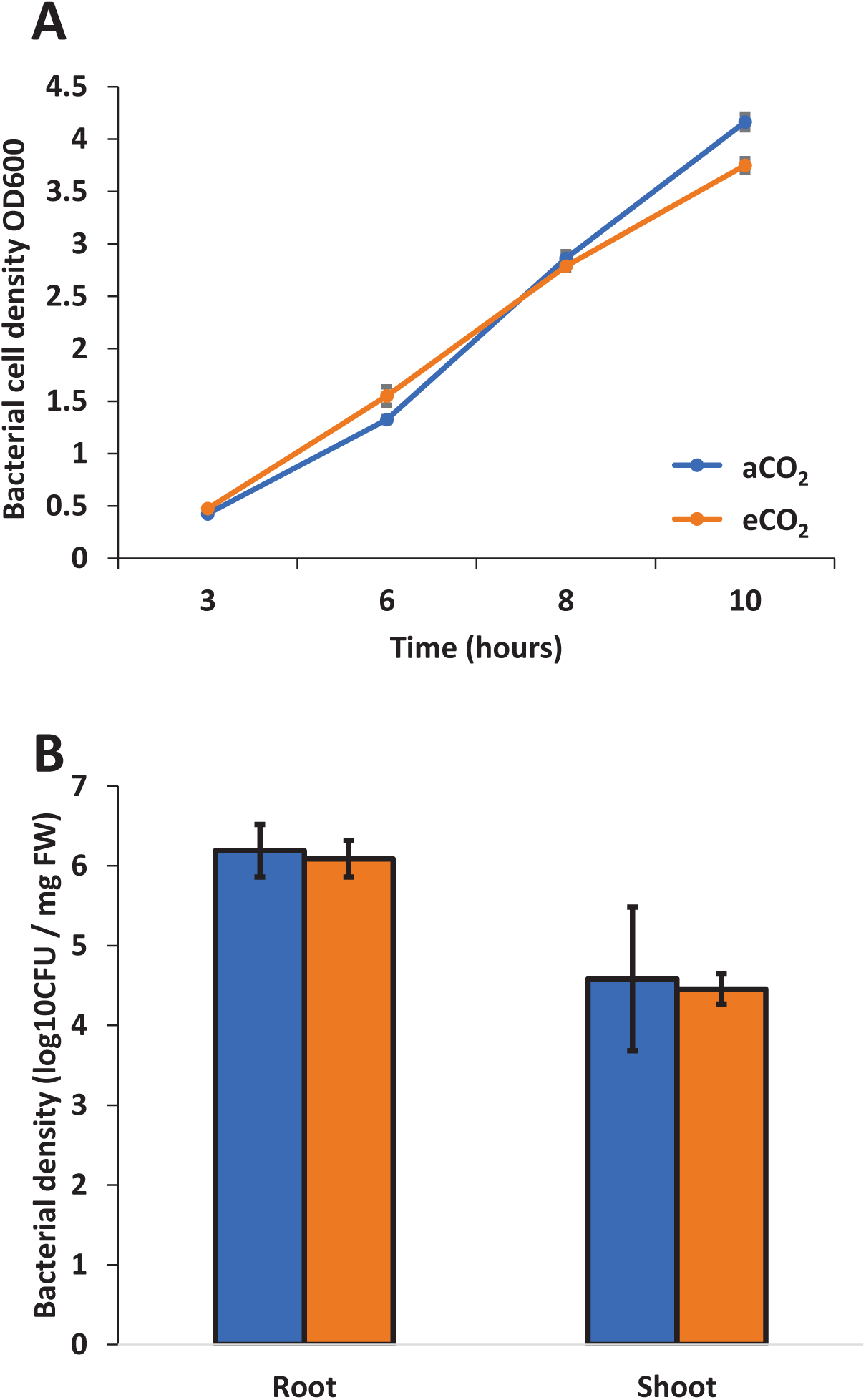
SA187 growth under aCO_2_ and eCO_2_ conditions. **(A)** SA187 growth in liquid LB culture under agitation at 30°C under aCO_2_ and eCO_2_ conditions, data represent the mean of 3 independent experiments **(B)** CFU counting of SA187 bacteria retrieved from root or shoot organs of *Arabidopsis thaliana* plants grown under ambient (aCO_2_) and elevated CO_2_ (eCO_2_) conditions. Blue bars indicate values from aCO_2_ and orange from eCO_2_ conditions. Data represent means (n =3 independent experiments) with standard deviations.

Further, bacterial colonization of Arabidopsis was quantified by measuring colony-forming units (CFU) per mg of plant tissue FW under both CO_2_ conditions. Consistent with prior studies (de Zélicourt *et al*., 2018), under aCO_2_, SA187 was found predominantly colonizing the roots rather than the shoots, with respective densities of 10^^6.1^ CFU/mg FW and 10^^4.5^ CFU/mg FW. Similar colonization densities were observed under eCO_2_, demonstrating that eCO_2_ did not affect SA187 abundance in/on plant tissues (Figure 2B).

These results confirm that the plant growth-promoting effects of SA187 in eCO_2_ conditions were not associated with an increased bacterial proliferation or colonization under eCO_2_ when compared to aCO_2_.

### C. SA187 inoculation alters phytohormone signaling pathway and primary metabolism under eCO_2_

To elucidate the molecular basis of the impact of SA187 on plant growth and development under eCO_2_, RNA sequencing (RNA-seq) was performed on Arabidopsis shoots and roots. The analysis revealed differential expression of 4048 genes in shoots and 1256 genes in roots compared to control (mock) plants under eCO_2_ (Table S1). To gain a comprehensive overview, transcriptomics data were organized using hierarchical clustering (Figure 3 and Figure S1) and analyzed for Gene Ontology (GO) enrichment.

**Figure 3:**
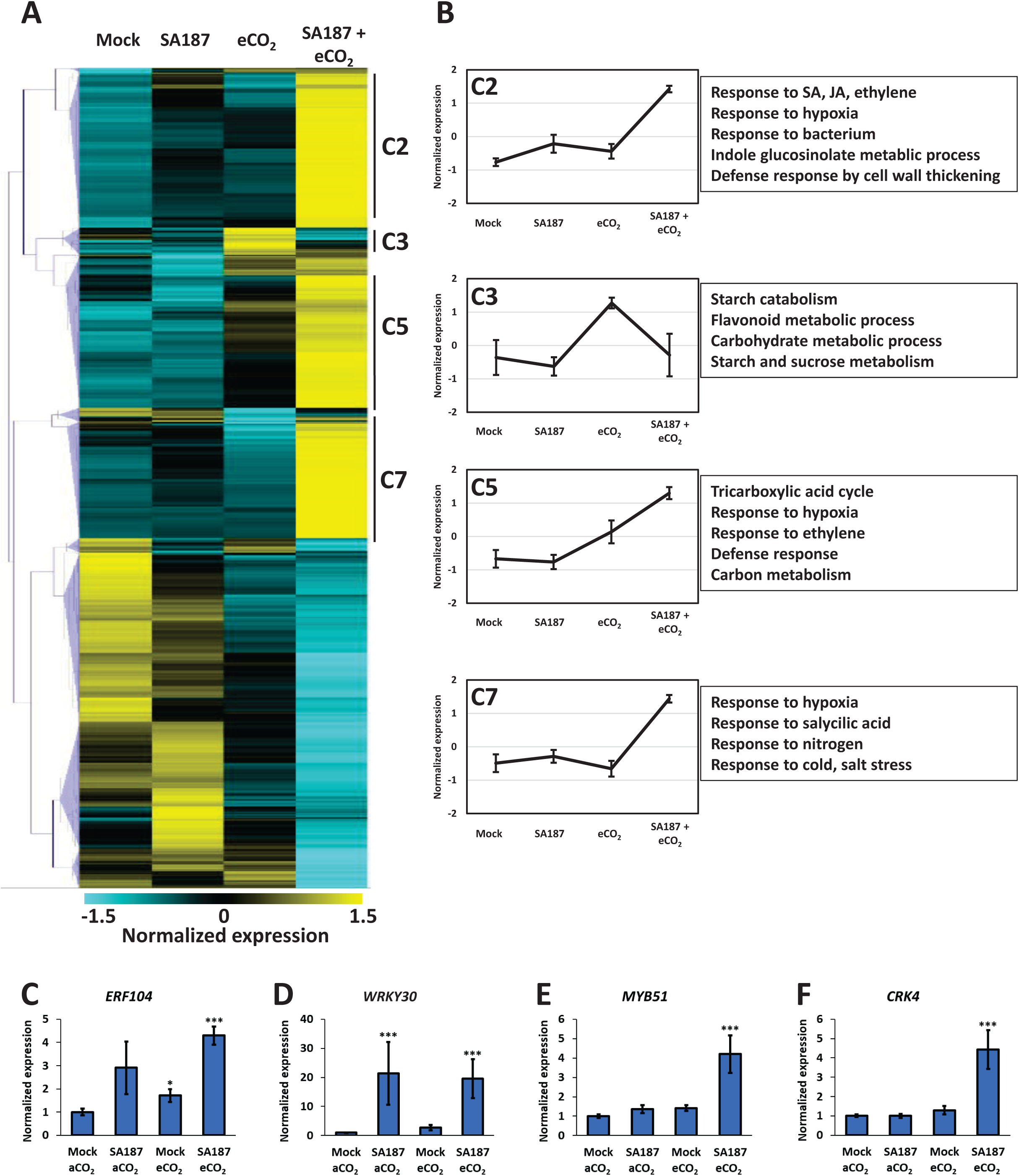
Arabidopsis shoot transcriptome analysis in response to SA187 inoculation and eCO2. **(A)** Heat map of DEGs in response to SA187, eCO_2_ both compared to mock plants grown in ambient conditions. Original mean counts were subjected to data adjustment by normalizing genes across all samples. Hierarchical clustering is displayed by average linkage under Pearson Correlation (MeV version 4). The colour scale indicates high and low expression levels. (**B)** Four selected clusters showing contrasting expression patterns between eCO_2_ and SA187 + eCO_2_ conditions using functional profiling with <u>G:Profile</u>r. qPCR expression analysis of **(C)** *ERF104*, **(D)** *WRKY30*, **(E)** *MYB51*, **(F)** *CRK4*, in shoots, in the four different tested conditions. Normalized expression indicates the linear fold change compared to mock-treated plants in aCO2. Data represent mean (n = 3 independent experiments) with standard errors. Asterisks indicate a statistical difference based on the Mann& Whitney test: \**p* < 0.05; \*\*\**p* < 0.001.

#### 1. Shoot Transcriptome

Hierarchical clustering grouped shoot DEGs into 11 clusters (Figure 3 and Sup Table 2). Clusters 2, 3, 5 and 7 appeared to be the most interesting, as they comprised DEGs regulated by SA187 under eCO_2_ (Figure 3). Overall, SA187 inoculation seemed to modify stress phytohormone homeostasis as GO terms associated with response to salicylic acid (SA), jasmonic acid (JA), and ethylene were induced in clusters 2, 5 and 7. Indeed, cluster 2 (772 genes) was comprised of genes that seemed to be unaffected by eCO_2_ alone but exhibited significant induction in response to SA187 under eCO_2_ conditions. This cluster was enriched in genes responsive to stress phytohormones SA, JA, and ethylene. Genes in Cluster 7 (580 genes) were similarly unresponsive to eCO_2_ alone but robustly induced upon SA187 inoculation under eCO_2_. Notably, in addition to SA responses, this cluster was also enriched in genes associated with N-metabolism. Genes in Cluster 5 (756 genes) exhibited a synergistic effect, being upregulated by eCO_2_ and further induced by SA187. Cluster 5 genes participate in primary metabolism, including the TCA cycle and C-metabolism, alongside defense responses and ethylene signalling (Figure 3B). In contrast to the previous clusters, genes in cluster 3 (136 genes) were highly induced by eCO_2_ alone but repressed by SA187 under eCO_2_, particularly those involved in carbohydrate metabolism (including starch and sucrose pathways).

#### 2. Root Transcriptome

In root tissues, five clusters were identified (Figure S1 and Sup Table 3). Cluster 5 was the largest cluster identified from the root RNA-seq data and contained 468 genes, that were highly induced by eCO_2_ and down-regulated by SA187 under both aCO_2_ and eCO_2_ conditions. This cluster was enriched in genes related to carbohydrate metabolic processes, but also in flavonoid biosynthesis and sulfur compound related processes (Figure S1). Conversely, cluster 2 included genes consistently upregulated by SA187, and independent of CO_2_ level. The genes in this cluster were enriched in defence responses and hypoxia, which is a typical SA187 response.

Taken together, the observed differential responses highlighted the role of SA187 in regulating shoot and root transcriptomes under eCO_2_. They indicated that the beneficial effect of SA187 was possibly mediated by the activation of stress phytohormonal signalling pathways. This was confirmed by RT-qPCR analyses of the ethylene marker *ERF104*, the SA marker *WRKY30*, and the stress and defense markers *MYB51* and *CRK4* (Figure 3 C-F). This induction was shown previously and ethylene was considered as the major signalling pathway to mediate SA187-dependent plant beneficial effects (de Zélicourt 2018, Shekhawat 2021). The transcriptomic analyses also revealed possible modifications of C (carbohydrates and starch, in cluster 3 and 5) and N metabolisms (cluster 7) that might help explain the improved plant growth under eCO_2_ when inoculated with SA187.

### D. SA187 Modulates TCA Related Metabolites and C/N Content Under eCO_2_

To complement the transcriptomic analyses, GC-MS-based metabolic profiling was carried out on extracts from Arabidopsis seedlings grown in the same 4 conditions in order to determine the metabolic changes occurring in plants when inoculated with SA187 in air and in eCO_2_. When compared to mock plants, out of the 114 metabolites identified and quantified, 20 metabolites were found to be differentially accumulated metabolites (DAM) in shoots in response to SA187 inoculation (Figure 4A and Sup Table 4), of which 9 were more abundant: beta-alanine, sinapic acid, tryptophan, trehalose, arbutin, nicotinic acid, oxoglutaric acid, pyruvic acid and asparagine. These metabolites are associated with diverse metabolic pathways including the TCA cycle (pyruvate and oxoglutarate), and amino acid metabolism (beta-alanine, asparagine and tryptophan) as well as defense responses (e.g. sinapic acid, arbutin). Regarding the reduced metabolites in response to SA187 inoculation, 11 of them were considered as less abundant, some being also associated to the TCA cycle such as citric acid, or amino acid with proline, or sugar with proline. This modulation of TCA related metabolites is partially in agreement with the transcriptome results, for instance the GO term TCA cycle was found to be significantly enriched in cluster 5, representing genes upregulated by SA187. In roots, 18 of 20 DAMs were significantly reduced including TCA cycle intermediates (succinic acid, fumaric acid) and amino acids such as beta-alanine, proline and glycine (Figure S2 and Sup Table 5).

**Figure 4:**
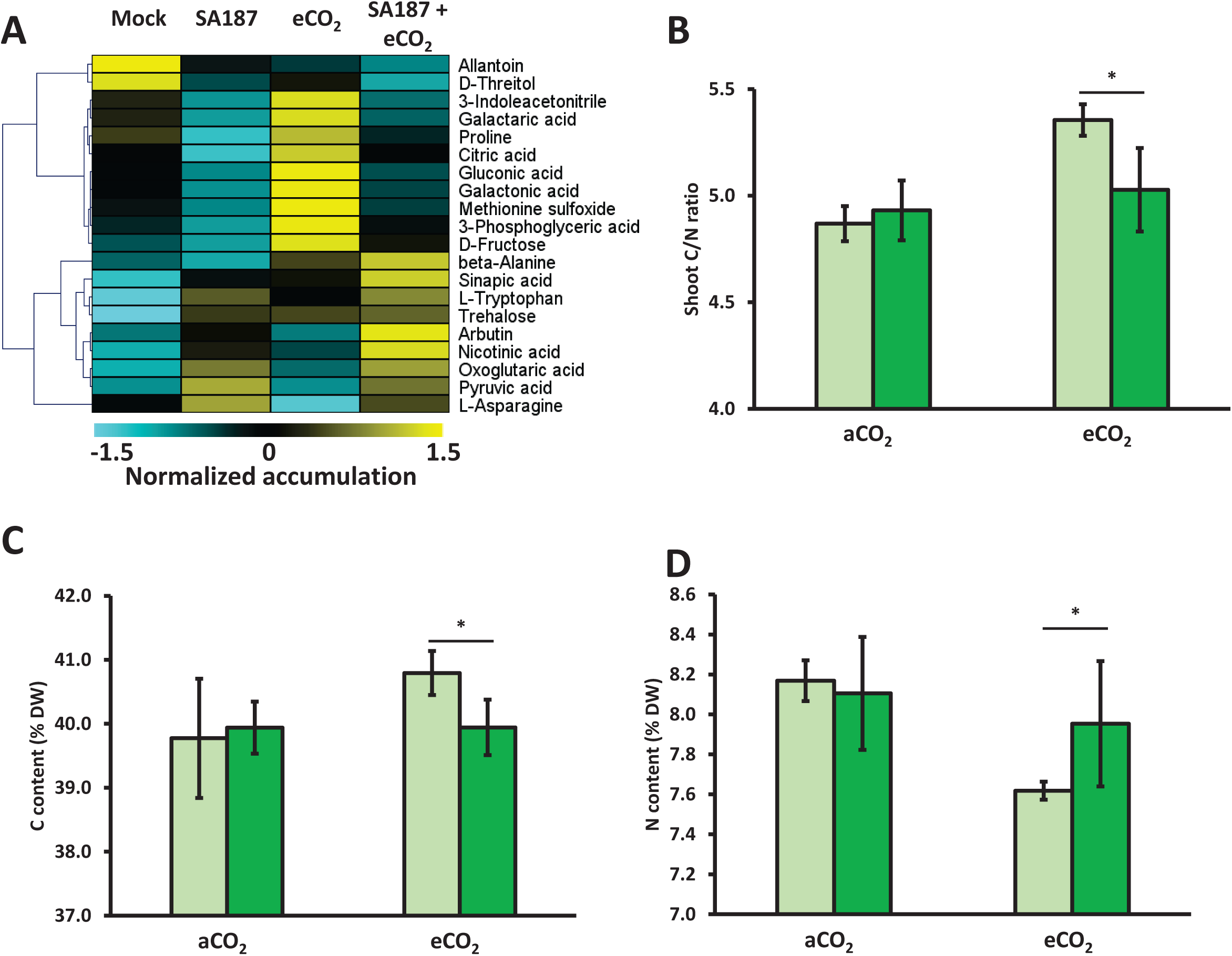
Arabidopsis shoot metabolic analysis in response to SA187 inoculation and eCO_2_. **(A)** Heat map of differentially accumulated metabolites in response to SA187 in either aCO_2_ or eCO_2_ conditions. Original mean counts were subjected to data adjustment by normalizing metabolites across all samples. Hierarchical clustering is displayed by average linkage under Pearson Correlation (MeV version 4). The colour scale indicates high and low accumulation levels. (**B)** Shoot C/N ratio, **(C)** shoot C content (% DW), **(D)** shoot N content (% DW) of mock (light green) and SA187 inoculated seedlings (dark green) under aCO_2_ or eCO_2_ conditions. Data represent means (n = 3 independent experiments) with standard deviations. Asterisks indicate a statistical difference based on the Mann & Whitney test: \**p* < 0.05.

Since the transcriptomic studies suggested that both C and N metabolisms could be deregulated in SA187 inoculated plants, elemental analyses were performed to quantify total C and N contents. Interestingly, no changes were observed for plant C, N contents and C/N ratio in response to SA187 inoculation in aCO_2_ conditions (Figure 4B-D). However, under eCO_2_ conditions whereas the C content was reduced upon SA187 inoculation, N content increased, thus leading to a significant decrease in plant C/N ratio to a value similar to that observed in aCO_2_.

Therefore, the metabolite profiling confirmed that SA187 inoculation modifies plant primary metabolisms including the TCA cycle and amino acid metabolism in agreement with the conclusions obtained from the transcriptomic analyses.

### E. SA187-induced Growth Promotion Under eCO_2_ Requires Ethylene Signalling

The transcriptomic analyses indicated a possible role for several phytohormones, among them ethylene, in the SA187-induced growth enhancement observed under eCO_2_ (Figure 3, cluster 2 and 5). To test whether ethylene signalling was involved, the SA187 response was compared between the ethylene-insensitive mutant *ein2-1* and wild-type Col-0 plants under eCO_2_ conditions.

As expected, SA187 significantly enhanced the FW of wild-type seedling under eCO_2_ (Figure 5A), consistent with previous observations (Figure 1). However, while an increase of PRL was still observed for *ein2-1* after SA187 inoculation (Sup Figure 3), seedling FW significantly decreased in SA187-inoculated *ein2-1* plants when compared to mock-treated *ein2-1* plants (−24% compared to mock conditions). This indicated that in the absence of a functional ethylene signalling pathway, SA187 shifted from promoting to inhibiting plant growth, and underscored that ethylene signalling was indispensable for the growth-promoting action of SA187 under eCO_2_, acting as a pivotal regulatory switch determining the bacterium’s influence on plant growth. This aligned with existing literature emphasizing ethylene’s key role in mediating plant responses to SA187 interactions and environmental stresses (de Zélicourt *et al*., 2018; Shekhawat *et al*., 2021).

**Figure 5:**
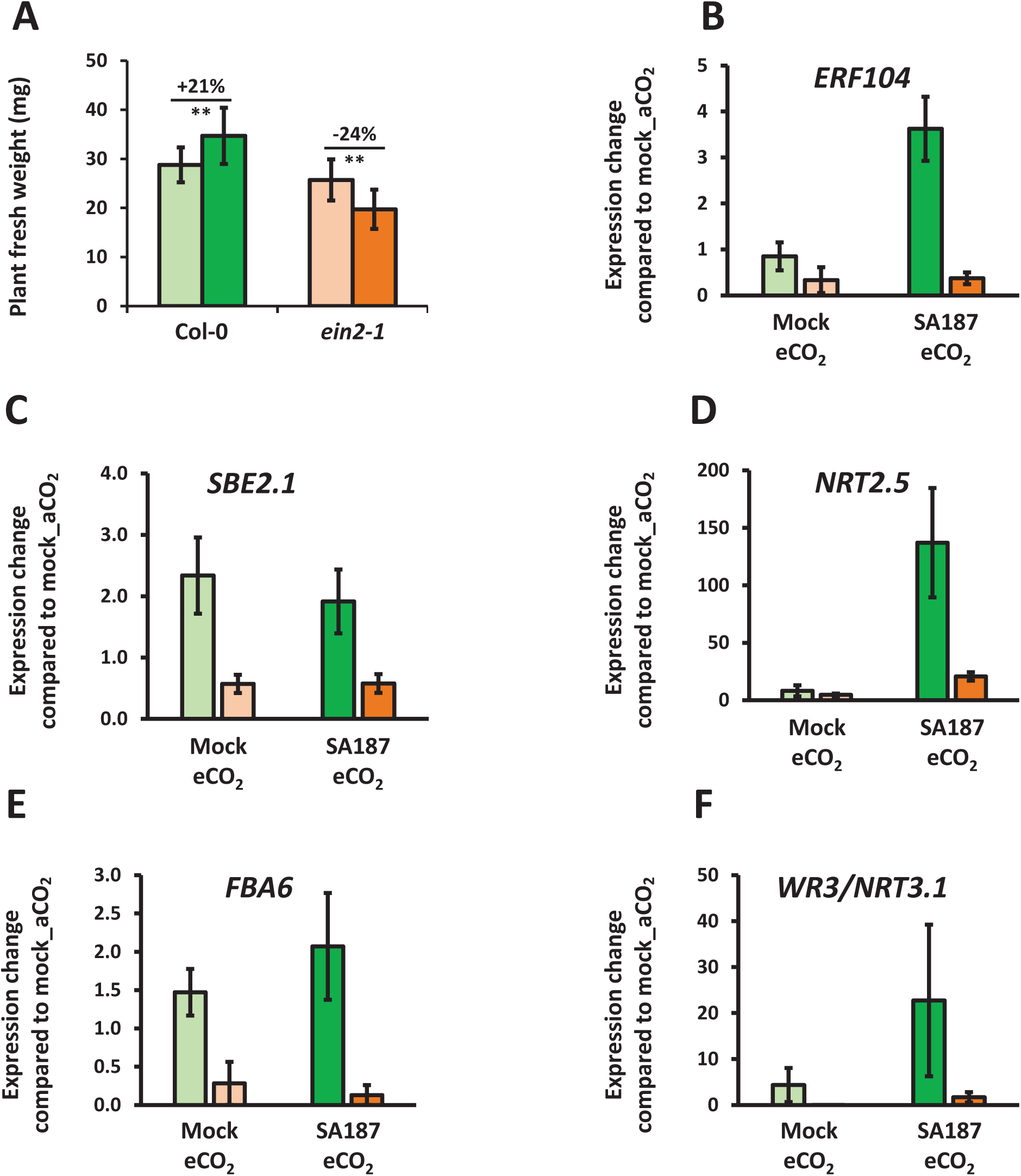
*ein2* response to SA187 inoculation under eCO2 conditions. **(A)** Plant fresh weight of mock and SA187 inoculated Col-0 and *ein2-1* plants in aCO_2_ and eCO_2_ conditions. qPCR expression analysis of **(B)** *ERF104*, **(C)** *SBE2.1*, **(D)** *NRT2.5*, **(E)** *FBA6* and **(F)** *WR3/NRT3.1* in Col-0 and *ein2-1* shoots under Mock (light green and light orange, respectively) or SA187 inoculation (light green and light orange, respectively) in eCO_2_ conditions. Expression values are indicated as linear fold changes compared to mock aCO_2_ conditions of a given genotype. Data represent means (n = 3 independent experiments) with standard errors. Asterisks indicate a statistical difference based on the Mann & Whitney test: \**p* < 0.05; \*\*\**p* < 0.001.

Given the transcriptomic insights into N et C metabolisms and ethylene signalling as key processes influenced by SA187 under eCO_2_, several genes representative of the GO terms significantly enriched in the transcriptomic datasets were selected to determine whether their regulation upon SA187 inoculation and/or eCO_2_ was still occurring in the absence of ethylene signalling: the ethylene marker gene *ethylene response factor 104* (*ERF104*, Cluster 7), *nitrate transporter 2.5* (*NRT2.5*, Cluster 7), *starch branching enzyme 2.1* (*SBE2.1*, cluster 3), *fructose-1,6-bisphosphate aldolase 6* (*FBA6*, Cluster 5), *wound responsive 3/nitrate transporter 3.1* (*WR3/NRT3.*1, Cluster 2). As expected *ERF104*, but also *NRT2.5*, and *WR3/NRT3.1* expression were no longer induced in response to SA187 in *ein2-1* when compared to wild type Col-0 (Figure 5B, D, F). Moreover, it appeared that *FBA6* and *SBE2.1*, associated to carbohydrate metabolism were unresponsive to both SA187 and to eCO_2_ conditions, as their expression were not induced (or even repressed) when compared to aCO_2_ conditions even in non-inoculated plants.

## IV. Discussion

Based on different models, the level of atmospheric CO_2_ is estimated to rise to 700-1000 ppm by the next century if no action is taken. Although eCO_2_ enhances photosynthetic CO_2_ assimilation and lower photorespiration rates, it has been shown that many C_3_ plants acclimate to long term eCO_2_, leading to a lower than predicted yield increase. This study aimed to determine whether inoculation with SA187 could increase plant growth and development under eCO₂ and to explore the underlying signalling and metabolic pathways.

### A. SA187 inoculation increases plant growth under eCO_2_

While SA187 inoculation had almost no effect on plant growth and development in aCO_2_ conditions, its significant effect on *Arabidopsis thaliana* under eCO_2_ suggested a synergistic interaction between bacterial inoculation and eCO_2_ levels. The observed increase in plant biomass under eCO_2_ conditions aligns with previous studies documenting the CO_2_ fertilization effect of C3 plants. Historically, this is primarily attributed to enhanced photosynthetic rates and a reduction of photorespiratory losses, together increasing net CO_2_ assimilation (Ainsworth & Long, 2005; Ainsworth and Long, 2020). Interestingly, SA187 led to an additional growth enhancement of both roots and shoots under eCO_2_ conditions when compared to mock-treated plants, suggesting that SA187 could modulate plant processes to optimize growth in high CO_2_ environments. Previous observations of a bacterial beneficial effect on plant growth under eCO_2_ conditions mainly concerned the symbiotic interaction between legume plants and N_2_-fixing rhizobacteria (Palit *et al*. 2020). That said, there are a few examples where a PGPR led to increased plant growth under eCO_2_. A good example concerns *Rahnella* sp. WP5, isolated from poplar, that maintained the photosynthetic activity of rice under eCO_2_ (Rho *et al*., 2019).

It should be noted that the growth-promoting effects of SA187 were independent of bacterial proliferation (Figure 2), suggesting that the observed phenotypic enhancements were due to intrinsic modifications within the plant in an eCO_2_ context, altering both metabolic and physiological processes.

Surprisingly, in our conditions, SA187 increased PRL, with no effect on LRD independently of atmospheric CO_2_ level. This feature contradicts previous findings indicating that SA187 did not affect PRL and only increased LRD under salt stress conditions (de Zélicourt *et al*., 2018). Although growth conditions were similar (culture medium, long-day conditions and temperature), differences could be due to site-specific growing conditions.

### B. SA187 Reprograms Plant Transcriptome and Modifies C & N metabolism

To obtain more insights into the molecular mechanisms affected by SA187 inoculation, transcriptomics and metabolic profiling were performed on separated roots and shoots of mock and SA187 inoculated plants under aCO_2_ and eCO_2_. Firstly, transcriptomic analysis revealed significant reprogramming of gene expression in both shoots and roots of Arabidopsis plants inoculated with SA187 under eCO_2_ conditions. GO term enrichment analyses identified significant enrichments for stress phytohormone signalling including genes responsive to SA, JA, and ethylene in shoots (Figure 3B). This is in agreement with previous reports showing that SA187 inoculation triggers such signalling pathways under normal conditions, but also in response to salt stress (de Zélicourt *et al*., 2018; Rolli *et al*., 2021). Induction of biotic stress responsive genes with no penalty on plant growth suggests that SA187 may enhance plant resistance to pathogens under eCO_2_, as SA and JA are well-known for their roles in plant defence responses, while ethylene is involved in various stress responses and developmental processes (Thilakarathne *et al*., 2025). This may be of a major importance in future climate change scenarios that are expected to facilitate plant disease outbreaks (Lahlali *et al*., 2024).

In addition to the induction of stress related phytohormones, it appeared that SA187 was also altering plant primary metabolisms in both shoots and roots. Bacterial colonization appeared to increase the expression of shoot TCA and carbohydrate metabolism related genes (Figure 3A, cluster 5) as exemplified by the induction of the glycolytic/gluconeogenesis gene *FBA6* encoding a fructose-1,6-bisphosphate aldolase (Figure 5). This was also supported by the accumulation of the final product of glycolysis and TCA precursor pyruvic acid, but also oxoglutaric acid (Figure 4A). On the other hand, starch metabolism gene expression appeared to be reduced, as observed in Shoot cluster 3, and *SBE2.1* gene expression analyses. Such observations suggest that inoculated plants invest more photosynthetically assimilated C into energy metabolism instead of C-reserves. Indeed, the TCA cycle is a central metabolic pathway that plays a crucial role in energy production and the biosynthesis of various organic acids associated with amino acid metabolism. Indeed, a stimulated TCA cycle activity could be related to the enhanced growth of the inoculated plants as overexpression of several TCA related enzymes have been shown to increase plant development and performance (Zhand & Fernie, 2023).

Additionally, N-metabolism also appeared to be stimulated in SA187 inoculated plants under eCO_2_ conditions as shown by the significant enrichment of genes associated with N-metabolism (cluster 7) and the induced expression of the high affinity nitrate transporter *NRT2.5* gene. It is possible that such changes in gene expression gave rise to an enhanced N-assimilation and metabolism that led to the observed increase in N-content and the altered C/N ratio under eCO_2_ conditions upon SA187 inoculation. This is particularly important as eCO_2_ is believed to lead to a N-dilution effect in plant tissues due to increased growth (Taub & Wang, 2008). By improving N-metabolism, SA187 may help maintain a balanced C/N ratio, which is an essential parameter to maintain optimal plant growth and development (Kant *et al*., 2012). This further supports the idea that SA187 could improve N-use-efficiency, a critical factor for sustaining the CO_2_ fertilization effect (Ruiz-Vera *et al*., 2017; Kant *et al*., 2012). Indeed, N is a key limiting nutrient for plant growth, and its efficient assimilation and metabolism would be essential for maximizing the benefits of eCO_2_ (Gojon *et al*., 2022; Rubio-Ascension and Bloom, 2017).

Concerning the root system, a lower number of genes were deregulated in the tested conditions resulting in a reduced number of clusters regrouping genes influenced by SA187 inoculation. One cluster (cluster 2) regrouped the genes induced by SA187 independently of CO_2_ levels, and their functions were associated with, among others, plant defence processes. This was not surprising as SA187 produces Microbe-Associated Molecular Patterns (MAMPs) that are recognized by the plant, which in return will activate defence mechanisms. This has been documented before, as the SA187 genome contains *fliC* genes encoding flagella proteins that have N-terminus sequences 60 to 80% similar to flg22 peptides, that will be recognized by the flg22 receptor FLS2 (Rolli *et al*., 2021). On the other hand, cluster 5 represented genes that were induced by eCO_2_ in mock plants compared to aCO_2_ mock plants and that SA187 inoculation appeared to alleviate. It regrouped genes associated to carbohydrate and sulphur compound metabolic processes and this is coherent with previously obtained results showing that SA187 inoculation reduced the expression level of sulphur related genes when grown under salt stress (Andres Barrao *et al*. 2021). The SA187-related reduction of carbohydrate metabolic processes deduced from transcriptomics appeared to be mirrored in SA187 inoculated roots including the reduced levels of TCA related intermediates such as succinic acid and fumaric acid. This and the reduced content in total C per mg DW might reflect an additional SA187 C-sink associated with the roots thus increasing sink strength from the shoot to the root. This scenario might lead to a reduced induction of starch related genes in the shoot of SA187 plants under eCO_2_. Indeed, both parameters have been shown to be important to explain plant acclimation to eCO_2_ (Jauregui *et al*., 2018; Ruiz-Vera *et al*., 2017, Krapp *et al*., 1999).

### C. Role of Ethylene Signalling

As genes involved in ethylene response were found to be significantly enriched in clusters 2 and 5 of the shoot transcriptomics (Figure 3), the requirement of this signalling pathway for SA187-induced growth promotion under eCO_2_ was tested using the ethylene-insensitive mutant *ein2-1*. The absence of growth enhancement in SA187 inoculated *ein2-1* underscores the major role of ethylene in mediating the beneficial effects of SA187. The lack of induction of SA187-responsive genes (*ERF104*, *NRT2.5*, *WR3/NRT3.1*) in *ein2-1* further supports the involvement of ethylene signalling in SA187-mediated growth promotion. Interestingly, the ethylene signalling pathway dependence of the increased expression of *NRT2.5* and *NRT3.1* further illustrates the complex interconnection between ethylene and N-response (Ma *et al*., 2022) and it is consistent with previous studies that have emphasised the pivotal role of ethylene in plant responses to microbial interactions and environmental stresses (de Zélicourt *et al*., 2018). Transcript analyses by qRT-PCR also showed the repression of genes associated with C and starch metabolisms (*FBA6*, *SBE2.1*) in *ein2-1* in response to eCO_2_ thus suggesting that ethylene signalling may also regulate primary metabolic pathways in response to eCO_2_ independently of SA187 inoculation. Our observations appear to agree with a previous report showing the importance of ethylene in plant adaptation to eCO_2_, indeed the *ein2-5* insensitive to ethylene mutant lacked the eCO_2_ fertilization effect and this correlated with a lack of activation of the starch associated genes (de Smet *et al*., 2020).

## V. Conclusions

This study supports SA187 as a potential biostimulant for improving crop performance under anticipated future atmospheric eCO_2_ levels. Interestingly the growth-promoting phenotypes of SA187 were independent of bacterial proliferation, as colonization levels did not vary significantly between aCO_2_ and eCO_2_ conditions. This suggested that the observed enhancements were due to intrinsic modifications within the plant in an eCO_2_ context, altering both metabolic and physiological processes. Transcriptomic, metabolic and physiological studies highlighted an extensive reprogramming of gene expression and altered metabolic processes in SA187-inoculated plants in eCO_2_ conditions. Major components included stress phytohormones like ethylene, and primary metabolisms such as the TCA cycle, amino acid and carbohydrate metabolisms. Such reprogramming appeared essential for the beneficial plant growth and development observed in an eCO₂ atmosphere, that may also enhance resilience to other environmental stresses. The contrasting responses observed in the *ein2-1* ethylene-insensitive mutant compared to wild-type plants emphasised the significance of ethylene signalling in mediating many of the observed SA187 effects.

Further work to better understand the molecular mechanisms by which SA187 influences plant responses under eCO_2_ would be useful. This could involve genetic, transcriptomic, and metabolomic studies using specific ethylene and N-uptake and signalling mutants. Metabolic flux analyses could offer deeper insights into how plant metabolism is rewired and C and N are dynamically allocated within SA187-inoculated plants under eCO_2_. Such studies would be paramount to help decipher the complex interactions between beneficial microbes and plants in adapting to climate change and pave the way to develop bio-based solutions to mitigate the negative impact of climate change on food security. Integrating microbial treatments with genetic improvements could provide a holistic approach to sustainable agriculture, promoting resilience and productivity in crops as global CO_2_ levels continue to rise.

## Author Contributions

A.I., M.H., and A.d.Z. designed the experiments, analyzed the data, and interpreted the results. A.I. performed all experiments except RNA-seq and metabolomics. C.M. conducted the metabolomics and elemental analyses and contributed to data interpretation. S.P. and C.P.-L.R. performed RNA-seq experiments and carried out initial analyses. J.B. genotyped the plant lines and provided technical support. A.d.Z. supervised the project and secured funding. A.I. wrote the first draft of the manuscript and A.d.Z. prepared the figures. A.d.Z. and M.H. revised the manuscript.

## Author Approvals

All authors have seen and approved the final version of the manuscript. The work described has not been published previously and is not accepted for publication elsewhere.

## Data Availability

NGS2020_09_SA187 RNA-Seq project was deposited to the Gene Expression Omnibus of the National Center of Biotechnology Information (Edgard R. *et al*. 2002): submission to GEO/NCBI in progress). All steps of the experiment, from growth conditions to bioinformatic analyses, were detailed in CATdb database (Gagnot S. *et al*. 2007): http://tools.ips2.u-psud.fr.fr/CATdb/; registered as NGS2020_09_SA187 according to the MINSEQE ‘minimum information about a high-throughput sequencing experiment’.

## Acknowledgements

Amina Ilyas was supported by a PhD scholarship from the French Ministry of Higher Education and Research (Ministère de l’Enseignement supérieur et de la Recherche). This work has benefited from a French State grant (Saclay Plant Sciences, reference n° ANR-17-EUR-0007, EUR SPS-GSR) under a France 2030 program (reference n° ANR-11-IDEX-0003). The authors would like to thank all students, colleagues and collaborators who contributed to discussions and technical support throughout the course of this study.

## List of supplemental information

**Supplemental Figure 1: Arabidopsis root transcriptome analysis in response to SA187 and eCO_2_.**

**(A)** Heat map of DEGs in response to SA187 in either aCO_2_ or eCO_2_ conditions. Original mean counts were subjected to data adjustment by normalizing genes across all samples. Hierarchical clustering is displayed by average linkage under Pearson Correlation (MeV version 4). The colour scale indicates high and low expression levels. **B** Two selected clusters showing contrasting expression patterns between eCO_2_ and SA187 + eCO_2_ conditions using functional profiling with <u>G:Profiler</u>.

**Supplemental Figure 2: Arabidopsis root metabolic analysis in response to SA187 inoculation**

**(A)** Heat map of differentially accumulated metabolites in roots in response to SA187 inoculation in aCO_2_ or eCO_2_ conditions. Original mean counts were subjected to data adjustment by normalizing metabolites across all samples. Hierarchical clustering is displayed by average linkage under Pearson Correlation (MeV version 4). The colour scale indicates high and low accumulation levels.

**Supplemental Figure 3: Growth phenotyping of Arabidopsis *Col-0* and *ein2* grown under elevated eCO_2_ conditions**

**(A)** Root Length, **(B)** Lateral root density, of eCO_2_ grown Col-0 and *ein2-1* plants 12 dpg without (Mock) and with (SA187) SA187. Light and dark green indicate Col-0 mock and SA187 plants respectively and light and dark orange indicate *ein-1* mock and SA187 plants respectively. For each measured parameter and every condition, data represent means (n > 30, 3 independent experiments) with standard deviations. Asterisks indicate a statistical difference based on two-sided, unpaired Student’s t-test: \*\**p* < 0.01.

**Supplemental Table 1:** List qRT-PCR Primer sets used in this study.

**Supplemental Table 2:** List of genes deregulated in shoots, at least, one condition compared to Mock aCO2 condition.

**Supplemental Table 3:** List of genes deregulated in roots, at least, one condition compared to Mock aCO2 condition.

**Supplemental Table 4:** List of metabolites identified and semi-quantified in shoot by GC-MS analysis.

**Supplemental Table 5:** List of metabolites identified and semi-quantified in root by GC-MS analysis.

