## Supplemental Figure 1 for "Enhancement of Arabidopsis growth by *Enterobacter* sp. SA187 under elevated CO_2_ is dependent on ethylene signalling activation and primary metabolism reprogramming"

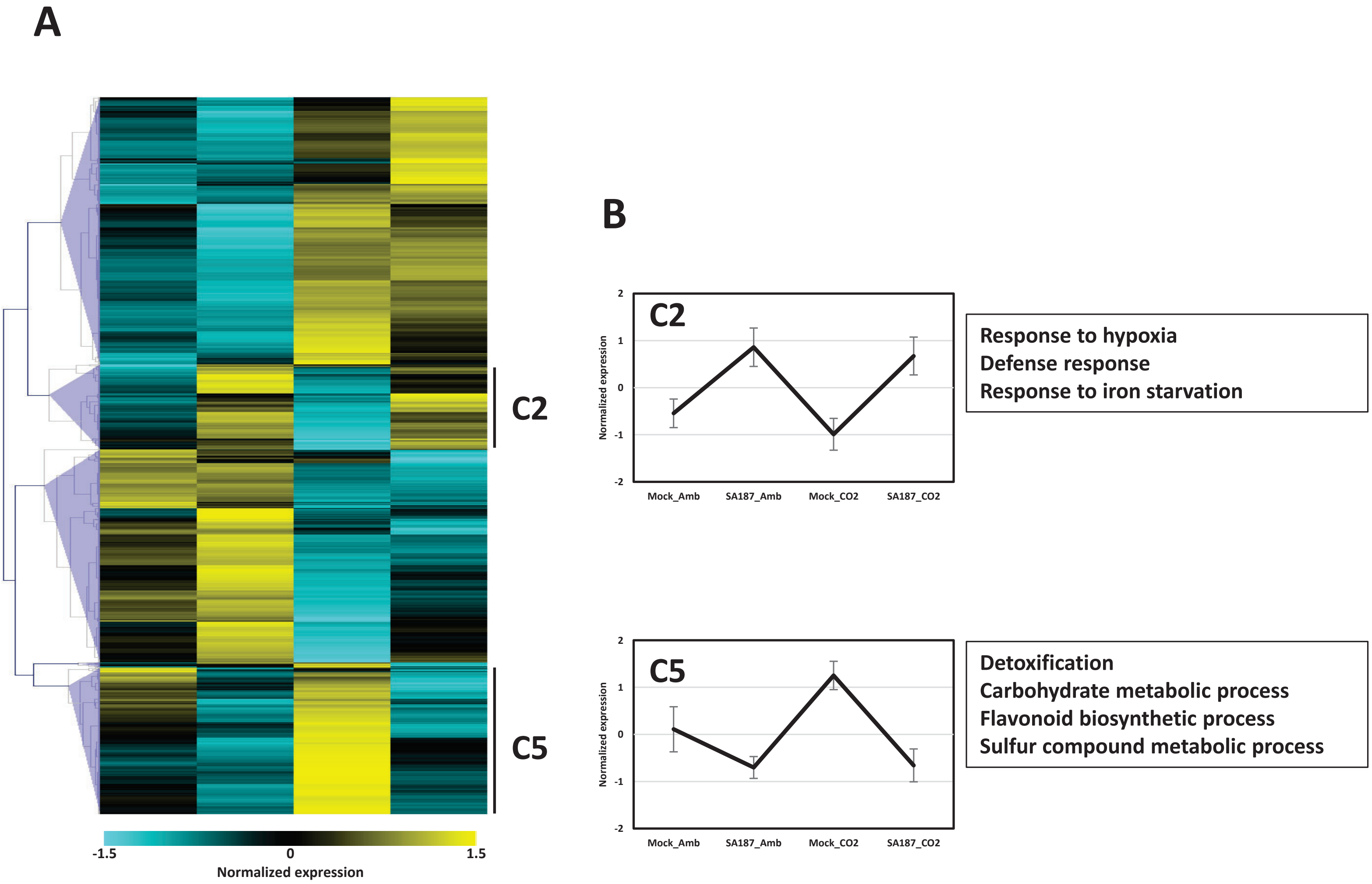


**Supplemental Figure 1: Arabidopsis root transcriptome analysis in response to SA187 and $eCO_2$.**
**(A)** Heat map of DEGs in response to SA187 in either $aCO_2$ or $eCO_2$ conditions. Original mean counts were subjected to data adjustment by normalizing genes across all samples. Hierarchical clustering is displayed by average linkage under Pearson Correlation (MeV version 4). The colour scale indicates high and low expression levels. **B** Two selected clusters showing contrasting expression patterns between $eCO_2$ and SA187 + $eCO_2$ conditions using functional profiling with G:Profiler.
