## Supplemental Figure 2 for "Enhancement of Arabidopsis growth by *Enterobacter* sp. SA187 under elevated CO_2_ is dependent on ethylene signalling activation and primary metabolism reprogramming"

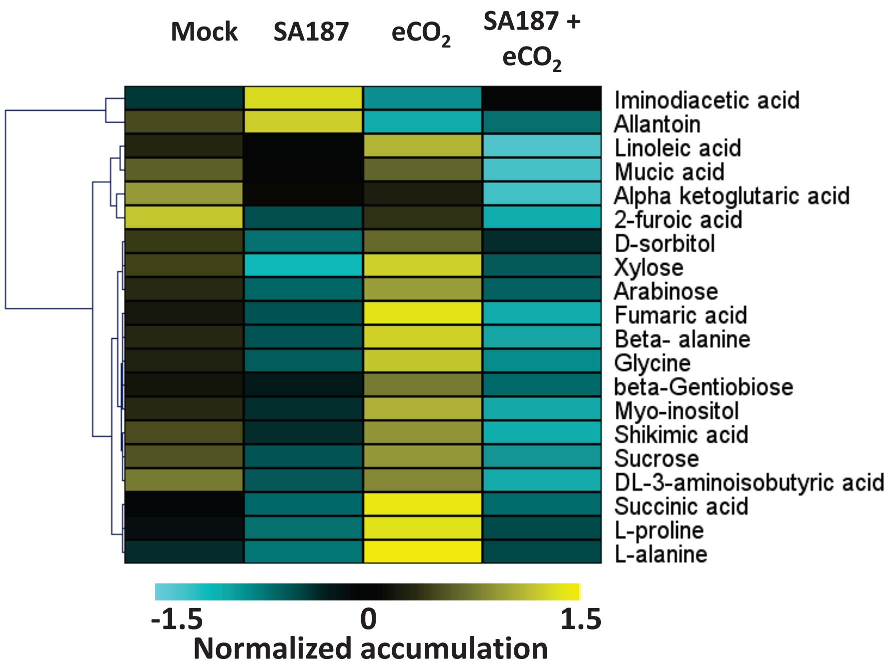


**Supplemental Figure 2: Arabidopsis root metabolic analysis in response to SA187 inoculation**

Heat map of differentially accumulated metabolites in roots in response to SA187 inoculation in $aCO_2$ or $eCO_2$ conditions. Original mean counts were subjected to data adjustment by normalizing metabolites across all samples. Hierarchical clustering is displayed by average linkage under Pearson Correlation (MeV version 4). The colour scale indicates high and low accumulation levels.
