## Supplemental Figure 3 for "Enhancement of Arabidopsis growth by *Enterobacter* sp. SA187 under elevated CO_2_ is dependent on ethylene signalling activation and primary metabolism reprogramming"

A

B

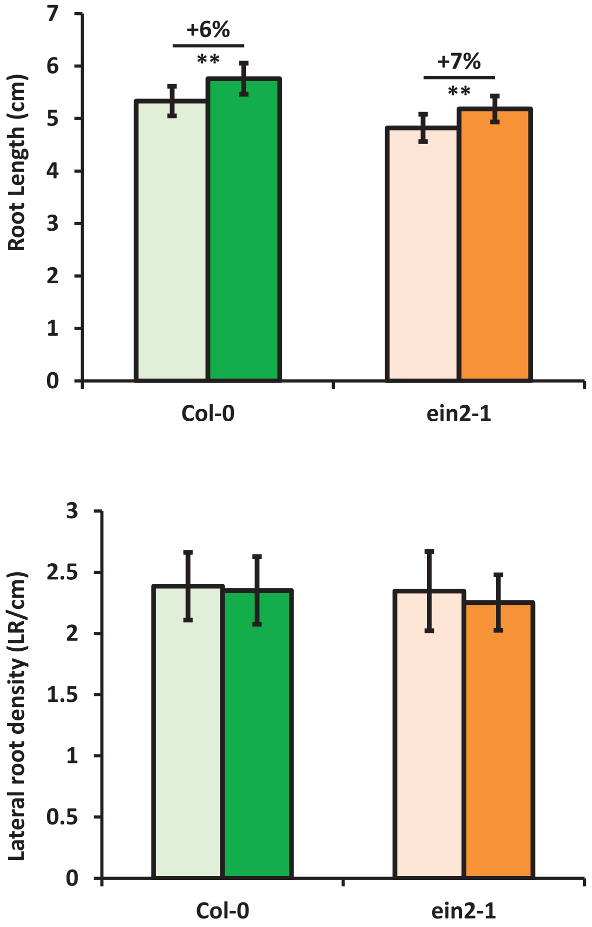


**Supplemental Figure 3: Growth phenotyping of Arabidopsis *Col-0* and *ein2* grown under elevated e$CO_2$ conditions**
**(A)** Root Length, **(B)** Lateral root density, of e$CO_2$ grown Col-0 and *ein2-1* plants 12 dpg without (Mock) and with (SA187) SA187. Light and dark green indicate Col-0 mock and SA187 plants respectively and light and dark orange indicate *ein-1* mock and SA187 plants respectively. For each measured parameter and every condition, data represent means (n > 30, 3 independent experiments) with standard deviations. Asterisks indicate a statistical difference based on two-sided, unpaired Student's t-test: ***p* < 0.01.
